# Primed-to-naive conversion of pluripotent stem cells can be tracked by specific DNA methylation changes for optimized culture conditions

**DOI:** 10.64898/2026.01.16.699914

**Authors:** Ian O. Shum, Thomas Akkermann, René Krüger, Kira Zeevaert, Wolfgang Wagner

## Abstract

During early embryonic development, cells transition from naive to primed pluripotent state. Various culture conditions have been established to revert primed cells back to naive state, to increase differentiation potential and to reset epigenetic abnormalities. In this study, we modified culture conditions to allow primed-to-naive conversion under feeder-independent and normoxic conditions (FINO medium), which exemplified the need for a quantitative measure of pluripotent states. DNA methylation (DNAm) profiling revealed extensive hypomethylation at naive state, but also significant gains of methylation at specific sites in the genome. We demonstrate that DNAm patterns can be used to benchmark culture protocols. Furthermore, we developed a naive-score based on DNAm at two genomic sites, which can be analyzed by digital PCR to monitor transition between pluripotent states. Our study describes a simplified culture protocol for primed-to-naive conversion, offers insights into the specific DNAm changes, and introduces a robust DNAm-based biomarker to track this process effectively.

## Introduction

Pluripotent stem cells (PSCs) exist along a spectrum of developmental states, ranging from the naive pre-implantation epiblast to the primed post-implantation epiblast (1, 2). Cultured naive PSCs are still capable to generate extra-embryonic lineages such as trophoblasts and amnion (3–5) and can efficiently form embryo-like blastoid structures (6–8). While murine PSC culture conditions are usually considered to provide naive state (9), conventional cultures of human PSCs more closely resemble the primed epiblast (10, 11). However, primed human PSCs often exhibit variable differentiation capabilities and tend to accumulate epigenetic abnormalities during long-term culture, which is particularly evident in genes affected by X chromosome inactivation (XCI) and genomic imprinting (12, 13). Notably, a transient pulse to the naive state during reprogramming has been demonstrated to effectively reset such abnormalities (14). This has spurred extensive efforts aimed at converting primed PSCs back into the naive state (15).

Over the past decade, numerous culture media have been developed to induce or stabilize naive-like pluripotency in human cells. These media utilize various combinations of inhibitors targeting pathways such as glycogen synthase kinase 3 beta (GSK3β), the mitogen-activated protein kinase (ERK1/2) pathway (16), the Wnt pathway (17), histone deacetylases (18), growth factors such as fibroblast growth factor, and other small molecule inhibitors (19, 20). A significant challenge of these protocols to support naive phenotype is their reliance on feeder layers. To this end, researchers identified a feeder-independent naive embryonic stem cell medium (FINE) through high-throughput chemical screening (21). Furthermore, most protocols depend on hypoxic conditions, which necessitates specialized incubators. While it has recently been shown that a naive-like state can be maintained under non-hypoxic conditions by a culture protocol utilizing the histone H3 methyltransferase disruptor of telomeric silencing 1-like inhibitor (DOT1L) along with a gamma-secretase inhibitor, this method still required feeder layers (22). Consequently, cultivating naive-like PSCs remains technically demanding. Moreover, it is still uncertain how closely these cultured cells mimic *in vivo* naive pluripotency, because cellular characteristics may change *in vitro* and quantitative comparisons are challenging (23).

So far, various indicators are used to validate naive phenotype, including dome-shaped morphology, single-cell survival rates, the presence of surface antigens, naive-specific transcription patterns, X-chromosome reactivation, differentiation potential into extraembryonic lineages, contribution to chimeras, and long-term genomic stability (24–27) – however, none of these criteria are well-suited as a unique biomarker for routine quality control or large-scale screening. Additionally, a global loss of DNA methylation (DNAm) is recognized as a hallmark of naive PSCs (18, 28). Yet, it has hardly been analyzed if specific CG dinucleotides (CpGs) undergo preferential hypomethylation during the transition to the naive state, and if other CpGs may even become hypermethylated in this process. DNAm patterns are widely used as biomarkers for other cell preparations (29, 30), because each cell type has characteristic epigenetic patterns, which are more stable than its transcriptome and ultimately determine the cellular identity (31). Furthermore, the intrinsic stability of DNA is advantageous for routine handling and DNAm can be interrogated quantitatively at single base resolution (32). Thus, the absence of specific DNAm biomarkers for classification of pluripotent states creates a significant gap in the current validation toolkit.

In this study, we sought to establish a feeder-independent, normoxic protocol (FINO) to ease primed-to-naive conversion. In fact, we observed that after five passages in FINO culture conditions the initially primed iPSC lines acquired typical characteristics of naive pluripotent cells, including dome-shaped morphology, up-regulation of relevant genes, and global hypomethylation – but this analysis also exemplified the need for a reliable and quantitative biomarker that can monitor the transition between the different states of pluripotency (33, 34). To this end, we anticipated that the primed-to-naive transition is not only reflected by global hypomethylation, but also by very specific gains and losses in the DNAm landscape. We therefore compiled DNAm profiles of 10 different studies and observed that across these studies specific CpGs revealed most drastic hypomethylation, whereas other sites even became consistently hypermethylated. Based on this, we established a targeted epigenetic signature of two CpGs – the naive-score. This score could reliably classify DNAm datasets into naive, primed, and intermediate state. Furthermore, we describe a time and cost-effective protocol to determine the naive-score by digital PCR to track transitioning between pluripotent states.

## Results

### Feeder-free and normoxic generation of naive-like stem cells

To ease primed-to-naive conversion we developed the FINO medium, which is a combination of the feeder-independent FINE formulation (21) with the non-hypoxia, LIF-dependent NHLD medium (22). In addition, we added ascorbic acid, which was previously used for naive culture conditions (35) as an antioxidant and cofactor for Ten-Eleven Translocation (TET) proteins (Figure 1a). Primed human iPSCs were switched to FINO culture conditions and monitored over several passages. The colonies started to grow in more compact clusters with the typical dome-shaped morphology of naive pluripotent cells (Figure 1b). This conversion was achieved repeatedly with at least four out of six iPSC lines tested, and subsequent experiments were performed with three independent iPSC lines. Gene expression analysis revealed a progressive up-regulation of naive markers (*DNMT3L*, *DPPA3*, *DPPA5*, *KLF5*, and *TFCP2L1*) alongside a concomitant down-regulation of primed genes (*HERVH*, *B3GAT1*, and *ZIC2*; Figure 1c). At passage 5, immunofluorescence confirmed strong nuclear KLF17 expression and cell-surface staining of Sushi domain-containing protein 2 (SUSD2), which are hallmarks of the naive state. Furthermore, we observed nuclear relocalization of hypoxia-inducible factor-1α (HIF-1α) even under normoxic conditions (Figure 1d, e). Flow cytometric analysis further validated the acquisition of naive surface antigens SUSD2 and CD75 with concurrent reduction of the primed marker CD90 (Figure 1f). Bulk RNA-seq identified 1024 up-regulated genes and 624 down-regulated genes in FINO-treated cells (adjusted p < 0.05, Log2 fold change > 1; Figure 1g). Gene set enrichment analysis of these differentially expressed genes highlighted significant positive enrichment (adjusted-*p* < 0.05) in mesenchymal epithelial transition and other pathways, including hypoxia-response, which is consistent with the observed nuclear HIF-1α localization (Figure 1h). Alternatively, we performed Gene Ontology analysis and observed significant enrichment of up-regulated genes in embryonic development categories, whereas down-regulated genes were rather associated with MHC class proteins (Supplemental Figure 1). Collectively, these data demonstrate that FINO medium efficiently drives the transition to a naive-like pluripotent state in a feeder-independent, normoxic environment.

**Figure 1.**
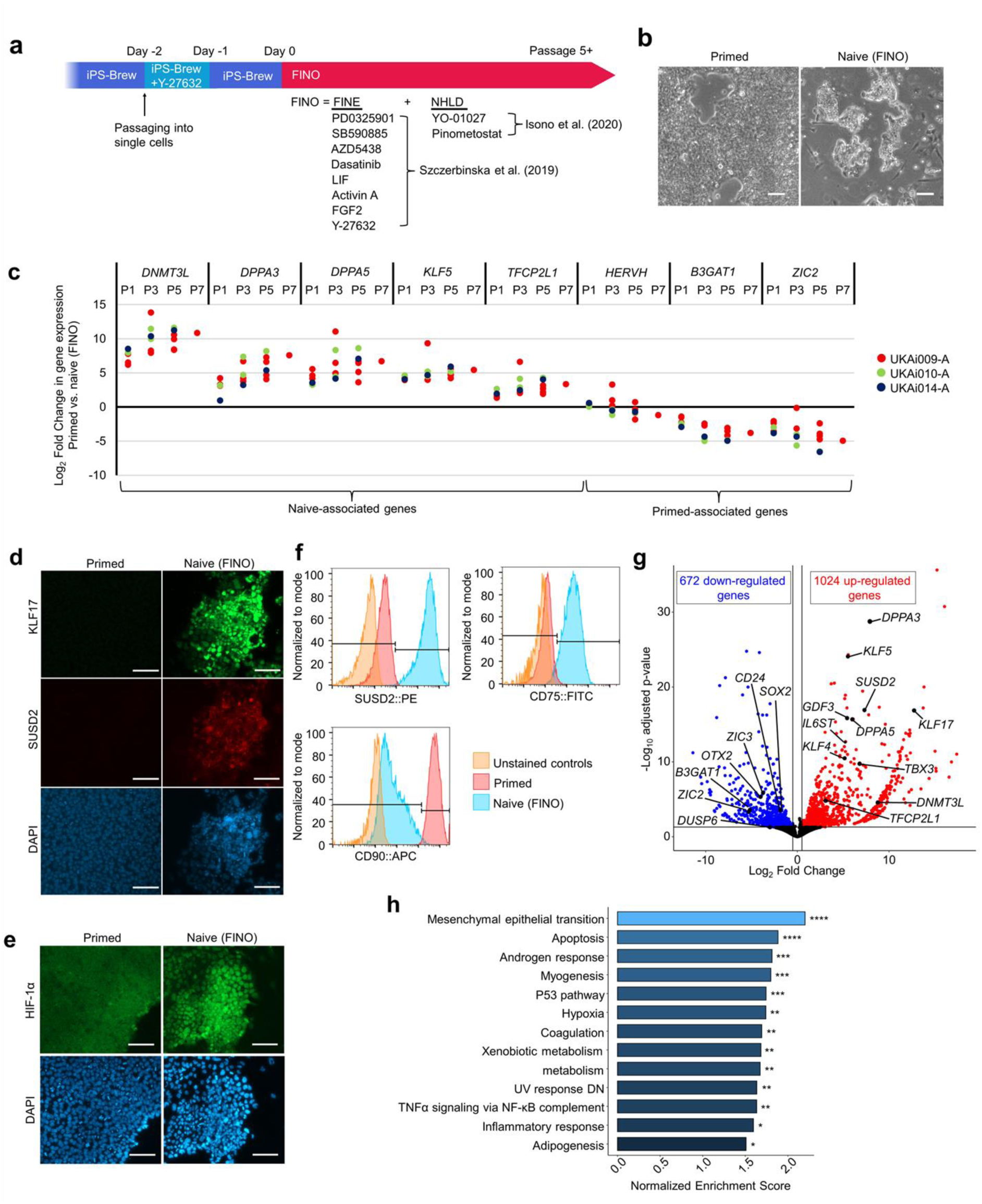
Primed-to-naive transition in a feeder-independent and normoxic environment. **(a)** Schematic presentation of the protocol and medium for feeder-independent normoxic conditions (FINO) for naive pluripotent stem cells. **(b)** Brightfield images of primed and FINO-treated pluripotent stem cells at passage 5. Scale bar = 100 µm. **(c)** Time course analysis of gene expression during FINO transition, showing upregulation of naive marker genes (*DNMT3L*, *DPPA3*, *DPPA5*, *KLF5,* and *TFCP2L1*) and downregulation of primed genes (*HERVH, B3GAT1*, and *ZIC2*). **(d)** Immunofluorescent images showing nuclear expression of KLF17 and cell surface staining for SUSD2 in FINO-treated cells. Scale bar = 100 µm. **(e)** Immunofluorescent images demonstrating nuclear relocalization of HIF-1α under normoxic conditions in FINO-treated cells at passage 6. Scale bar = 100 µm. **(f)** Flow cytometric analysis confirming the acquisition of naive surface antigens SUSD2 and CD75 with concurrent reduction of the primed marker CD90 in FINO-treated cells (passage 5). **(g)** Volcano plot showing 1024 up-regulated and 624 down-regulated genes upon FINO-treatment (passage 5; adjusted p-value < 0.05, Log2 fold change > 1). **(h)** Gene set enrichment analysis of the differentially expressed genes (adjusted p-value *< 0.05, **< 0.01, ***< 0.001, ****< 0.0001).

### DNA methylation profiles of FINO-treated cells

To determine whether FINO treatment evokes global DNA hypomethylation, another hallmark characteristic of naive pluripotent stem cells, we performed DNAm profiling. FINO treated cells displayed a pronounced shift toward lower methylation levels compared to primed cells (Figure 2a). Differential methylation analysis identified 210,529 CpG sites (32.38% of CpGs analyzed) as significantly hypomethylated (adjusted-*p* value < 0.05, mean difference in DNAm > 0.2). To our surprise, we also observed significant gain of DNAm at 8,333 CpG sites (1.28%), indicating that the remodeling of the methylome does not only involve hypomethylation (Figure 2b). When we examined DNAm levels at CpGs within the 1 kb flanking region of these significant hyper-(Figure 2c) or hypomethylated sites (Figure 2d), we observed that overall neighboring CpGs showed a similar trend of differential methylation within the 2 kb window. The hypomethylated CpGs showed a significant enrichment at CpG islands and particularly at the surrounding shore regions, whereas this was not observed for the hypermethylated regions (Figure 2e). Furthermore, hyper-and hypomethylated CpGs were rather enriched at intergenic regions (Figure 2f). We exemplarily focused on DNAm at CpGs in the naive-associated genes *DNMT3L, DPPA3*, *DPPA5*, *TFCP2L1*, and *KLF5*, which revealed pronounced hypomethylation at most of the corresponding CpGs (Figure 2g), whereas this was less consistent for genes that are upregulated at primed state or during differentiation (Supplemental Figure 2). When we matched the DNAm changes in promoter regions with corresponding gene expression, we observed in tendency an inverse correlation (Figure 2h). X-chromosome inactivation is reflected by DNAm, and hence transcriptional silencing of the inactive X chromosome. In fact, in our two female iPSC lines this is clearly reflected in the methylome at primed state: When we focused on CpGs on the X-chromosome, which were excluded for the rest of our study, we observed two peaks of DNAm levels around 100% and 50% (Figure 2i). Thus, at least one of the two X-chromosomes was always methylated due to XCI, whereas this was not observed in all other chromosomes (Figure 2a). Notably, in FINO treated cells the 50% DNAm level peak disappeared, and CpGs on the X-chromosome became overall demethylated, which indicates X-chromosome reactivation at naive state.

**Figure 2.**
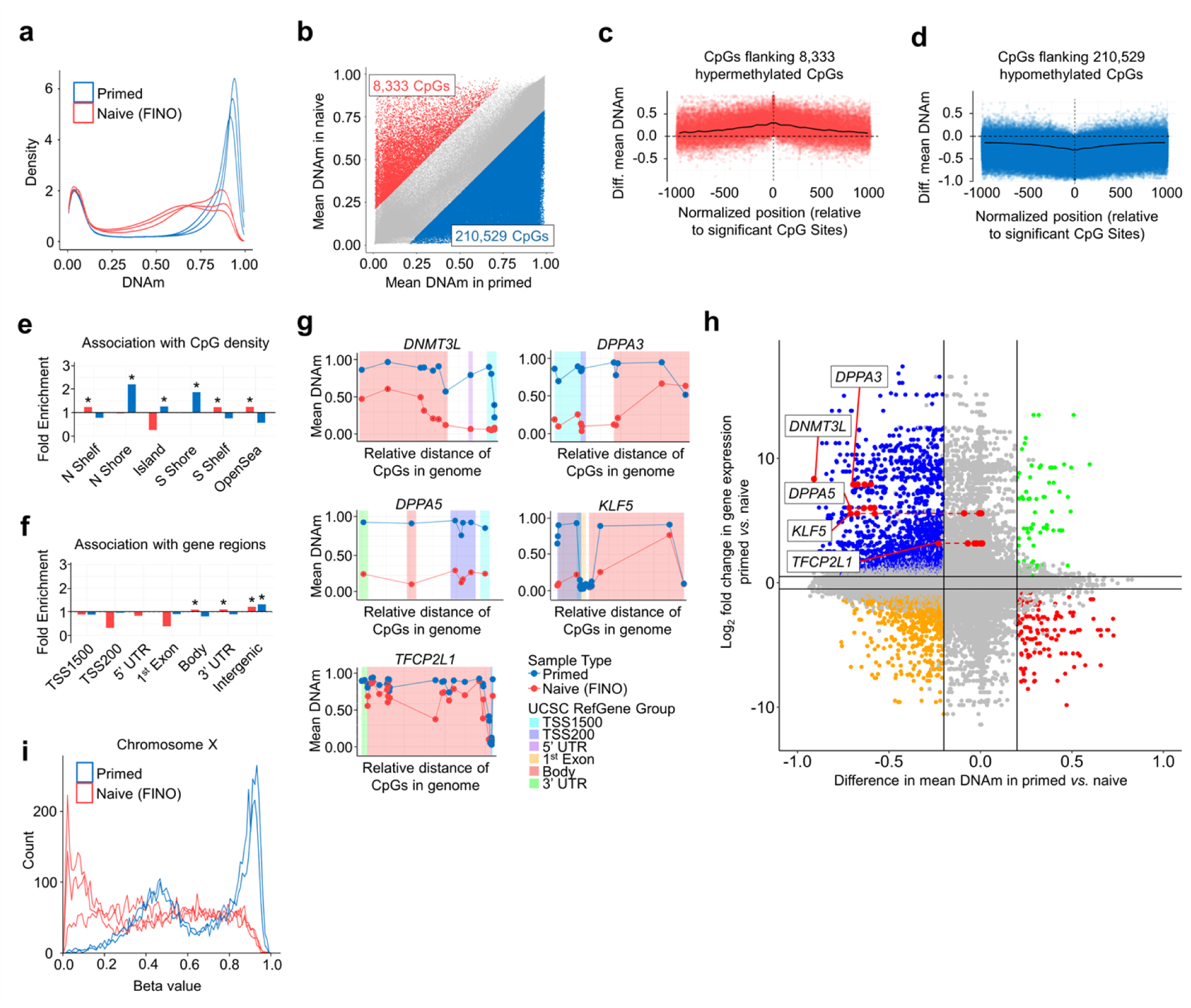
Epigenetic changes in primed-to-naive conversion with the FINO protocol. **(a)** Histograms of global DNA methylation levels in primed and FINO-treated cells (X and Y chromosomes excluded). **(b)** Scatter plot of differential methylated CpGs, with 210,529 hypomethylated CpG sites and 8,333 hypermethylated sites in FINO *versus* primed cells (adjusted p value: < 0.05, diff. mean DNAm > 0.2). **(c-d)** Mean DNAm changes within a 1kb adjacent to all significant hyper-and hypomethylated CpGs. **(e-f)** Enrichment analysis of significant hyper-and hypomethylated CpGs in primed stem cells and FINO-treated cells in relation to (e) CpG islands and (f) gene regions. (Hypergeometric distribution: * *p* < 0.05). **(g)** DNAm values at CpGs associated with five representative marker genes for naive state (*DNMT3A*, *DPPA3*, *KLF5*, *TFCP2L1*, and *DPPA5*) showing nearly all promoters (TSS1500 and TSS200) and gene bodies (1^st^ Exon and Body) become hypomethylated under FINO conditions. **(h)** Scatter plot of mean difference in DNAm *versus* fold-change in RNA expression (both primed/FINO-treated cells). Only CpGs in promotor regions are depicted (each gene can be associated with several CpGs and the five candidate genes are highlighted; colors depict significance in both comparisons: adjusted p-value < 0.05, fold-change gene expression > 2, mean difference in DNAm > 0.2). **(i)** Histograms of DNAm levels of CpGs on the X-chromosome in primed and FINO-treated cells.

### Specific modulation of the DNAm landscape between pluripotent cell states

Based on our findings that specific CpGs gain or lose DNAm in primed-to-naive conversion, we investigated how the DNAm landscape changes across different established protocols. We curated ten public DNAm datasets (dataset 1–10; Supplemental Table S1) that utilized a wide range of culture media for primed-to-naive conversion, alongside our primed and FINO samples (dataset 11). Multidimensional scaling of global DNAm patterns showed that primed and naive cell types clustered across the different datasets – our FINO-treated cells clearly clustered with other naive culture conditions, whereas the datasets 1-5 remained closer to the primed cells (Figure 3a). Next, we performed differential DNAm analysis for each of the datasets (adjusted *p*-value < 0.05, mean difference in DNAm > 0.2; Supplemental Figure 3a-j). As expected, all datasets showed a high number of significantly hypomethylated CpGs, but we also observed in all of them significant gains of DNAm in naive state. Notably, a high fraction of these gains and losses are overlapping between the different protocols in pairwise comparisons (Figure 3b). This indicated that despite the use of different culture conditions, overall similar DNAm changes were evoked in primed-to-naive conversion.

**Figure 3.**
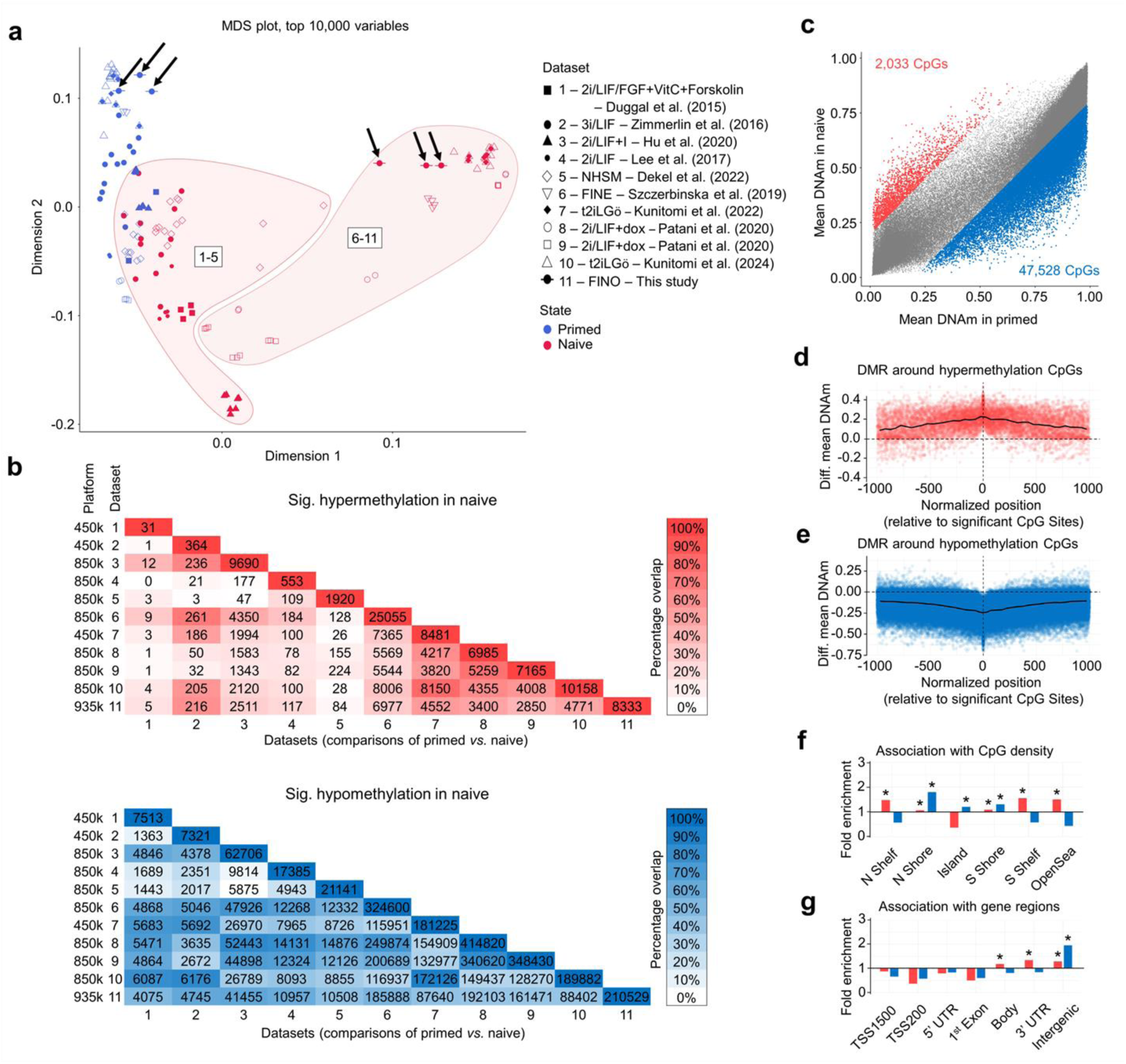
Integrative DNA methylation analysis of primed and naive stem cells. **(a)** We curated ten DNAm datasets on primed and naive pluripotent stem cells (dataset 1–10; all Illumina BeadChip data; Supplemental Table S1) to compare our profiles of primed and FINO-treated cells (dataset 11; highlighted by arrows). The multidimensional scaling (MDS plot) clustered DNAm profiles into primed and naive cells, albeit the studies 1-5 clustered closer to the primed state. **(b)** In each of the datasets, we determined the number of significantly hypo-and hypermethylated CpGs (adjusted *p*-value < 0.05, mean difference in DNAm > 0.2; Supplemental Figure S3), and the overlap of these CpGs is depicted for pairwise comparison between the datasets. **(c)** Scatter plot of differentially methylated CpGs in combined datasets 1-10 (adjusted *p*-value: < 0.05, mean difference in DNAm > 0.2). **(d,e)** Differential mean DNAm of CpGs within a 2kb window around significant (d) hyper-and (e) hypomethylated CpGs. **(f,g)** Enrichment analysis of significant hyper-and hypomethylated CpGs in datasets 1-10 in relation to (f) CpG islands and (g) gene regions (hypergeometric distribution: * *p* < 0.05).

We subsequently combined all ten public datasets to identify 47,528 hypomethylated CpGs (17.75% of CpGs analyzed), and 2,033 hypermethylated CpGs (0.75%; adjusted *p*-value < 0.05, mean difference in DNAm > 0.2; Figure 3c). Overall, these CpGs revealed very similar association with CpG islands (Figure 3d, e) and gene regions (Figure 3f, g), as mentioned above for our FINO-treated cells. However, there was no significant enrichment of these CpGs in Gene Ontology categories, which might either be attributed to the fact that the majority of differentially methylated CpGs were not in promotor regions, or to the heterogeneity between studies. Given that we saw closer clustering of our FINO cells with naive cells from datasets 6-10, we alternatively focused on these datasets and found 1,890 overlapping hypermethylated CpGs. When we analyzed the corresponding associated genes, we observed up-regulation of genes involved in early developmental patterning and down-regulated neuronal genes (Supplemental Figure 3k). Overall, these results clearly demonstrate that the transition between primed and naive pluripotent state is associated with very reproducible DNAm changes at specific sites in the genome – rather than just a global hypomethylation.

### Epigenetic biomarker for naive pluripotent stem cells

Given that the primed-to-naive transition is associated with particular changes in the DNAm landscape, we sought to establish an epigenetic signature that could serve as a biomarker for tracking this process. Our goal was to develop a targeted signature based on a few CpGs to facilitate easy and cost-effective applicability with different DNAm analysis methods. To identify these key CpGs, we used the published primed-to-naive transitions datasets (datasets 1-10) and utilized the CimpleG framework (36) to select the top six candidates, prioritizing those with significant mean differential DNAm and low intra-group variance (Figure 4a). Furthermore, we reasoned that our biomarker should not be affected by DNAm changes related to early cell fate decisions toward different germ layers. Therefore, we conducted a second selection using CimpleG against a reference set comprising primed and early differentiated cells directed toward mesoderm, endoderm, and ectoderm (37) (Figure 4b). This analysis revealed three CpGs that appeared particularly well-suited for characterizing the naive state: cg02351655, situated in intron 7 of the Coronin 7 gene (*CORO7*; hypermethylated in naive), cg19205850 in intron 1 of Ras-related protein Rab-7a (*RAB7A*), and cg25262261 in the promoter of Nuclear Cap Binding Protein Subunit 2 (*NCBP2*; both hypomethylated in naive; Figure 4c). None of these genes are associated with canonical pluripotency markers and they did not reveal significant differential expression in gene expression data (Supplemental Figure 4a-c).

**Figure 4.**
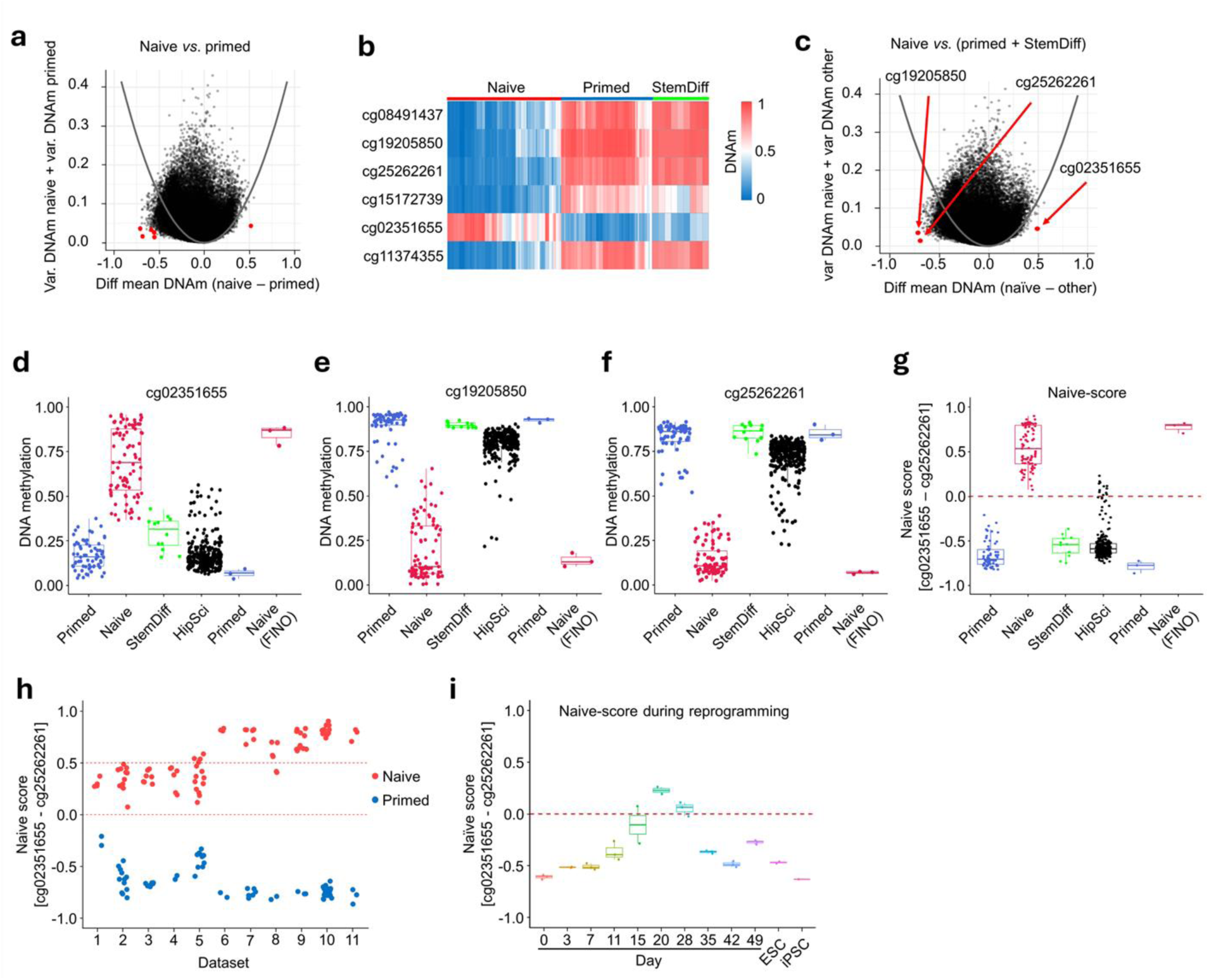
DNA methylation changes are consistent upon primed-to-naive transition. **(a)** CimpleG plot (36) of the selection of candidate CpGs to discern primed and naive DNAm profiles of datasets 1-10. The selection is based on the mean DNAm differences of primed and naive datasets and the variation within each of these groups. **(b)** Heatmap of DNAm levels of the six top CpGs in naive, primed, and early differentiation towards the three germ layers (StemDiff) (37). **(c)** Alternatively, we used CimpleG to select CpGs that could best discriminate naive DNAm profiles from primed and differentiated cells. **(d-f)** DNAm levels at the selected CpGs in the samples of primed and naive stem cells from dataset 1-10, early differentiated cells (StemDiff), primed stem cells from the iPSC cell bank (HipSci), and primed and naive stem cells from this study. **(g)** Naive-score (DNAm at cg02351655 – DNAm at cg25262261) for each of these samples. The dotted line depicts the arbitrary threshold at 0. **(h)** Naive-score for samples separated by datasets 1-11. The dotted lines depict thresholds for intermediate naive state (between 0 and 0.5). **(i)** Naive-score in DNAm profiles during reprogramming of somatic into pluripotent cells (42). At day 20 the cells transiently surpass the naive threshold.

We benchmarked these CpGs on the 66 primed and 85 naive samples of datasets 1 to 10, on the early differentiated cells toward the three germ layers (37) and on 275 iPSC samples from the HipSci stem cell bank, which were cultured under primed culture conditions (38), as well as our primed and FINO-treated cells (Figure 4d–f). All three CpGs could discern primed and naive samples, while only few samples of the HipSci showed DNAm levels indicative for naive state. Since all three CpGs showed similar results, we opted combining a hyper and a hypomethylated CpG into a “naive-score” to further ease applicability (naive-score = DNAm [cg02351655] – DNAm [cg25262261]). As the hypomethylated site, we chose cg25262261 because it had slightly higher discriminatory power than cg19205850. The threshold to classify into primed or naive cell preparations was arbitrarily set at 0. Notably, this analysis correctly classified all primed, all naive, and all early differentiated samples with 95.63% of the cell lines from the HipSci database being correctly classified as primed (Figure 4g). The few HipSci samples with higher naive-score showed consistent naive-like patterns at all three CpGs, indicating that these samples might in fact resemble a more naive-like methylome (Supplemental Fig 4d-f).

There is evidence that the various different protocols for primed-to-naive conversion result in a spectrum of naive states, and particularly the t2iLGö condition (dataset 10) is considered to closely resemble the pre-implantation naive epiblast state (39). In fact, when we compared the naive-score in the individual datasets, we observed that datasets 1 to 5 resulted in a lower naive-score than the datasets 6 to 11, indicating that they might be classified as “intermediate naive” (threshold for the score arbitrarily set between 0 and 0.5; Figure 4h).

Subsequently, we investigated how the naive-score changes during reprogramming of somatic cells into iPSCs. Various studies demonstrated that the cells acquire a naive-like dome-shaped morphology in this process (39, 40) and single cell transcriptomic analysis also suggested coexistence of primed-like and naive-like cells during reprogramming (41). Notably, when we analyzed the naive-score in a dataset that monitors DNAm changes during reprogramming (GSE54848) (42), we observed a transient state surpassing of the naive threshold at day 20, followed by a rapid loss of naive epigenetic character (Figure 4i; Supplemental Fig. 4g-i).

### Tracking of pluripotent states by digital PCR

To develop an easily applicable and cost-effective assay for the naive-score, we designed digital PCR assays to determine the DNAm levels at the relevant CpGs (Supplemental Figure 5a-c). Our protocol could demarcate methylated and non-methylated DNA strands within six hours (Figure 5a) to provide DNAm levels that closely resembled the measurements from the Illumina BeadChips (Supplemental Figure 5d). When we tested the dynamics during primed-to-naive conversion in FINO medium for independent cell preparations, all three tested CpGs showed rapid DNAm changes even within the initial passages (Figure 5b-d), and after five passages the naive-score reached a plateau (Figure 5e).

**Figure 5.**
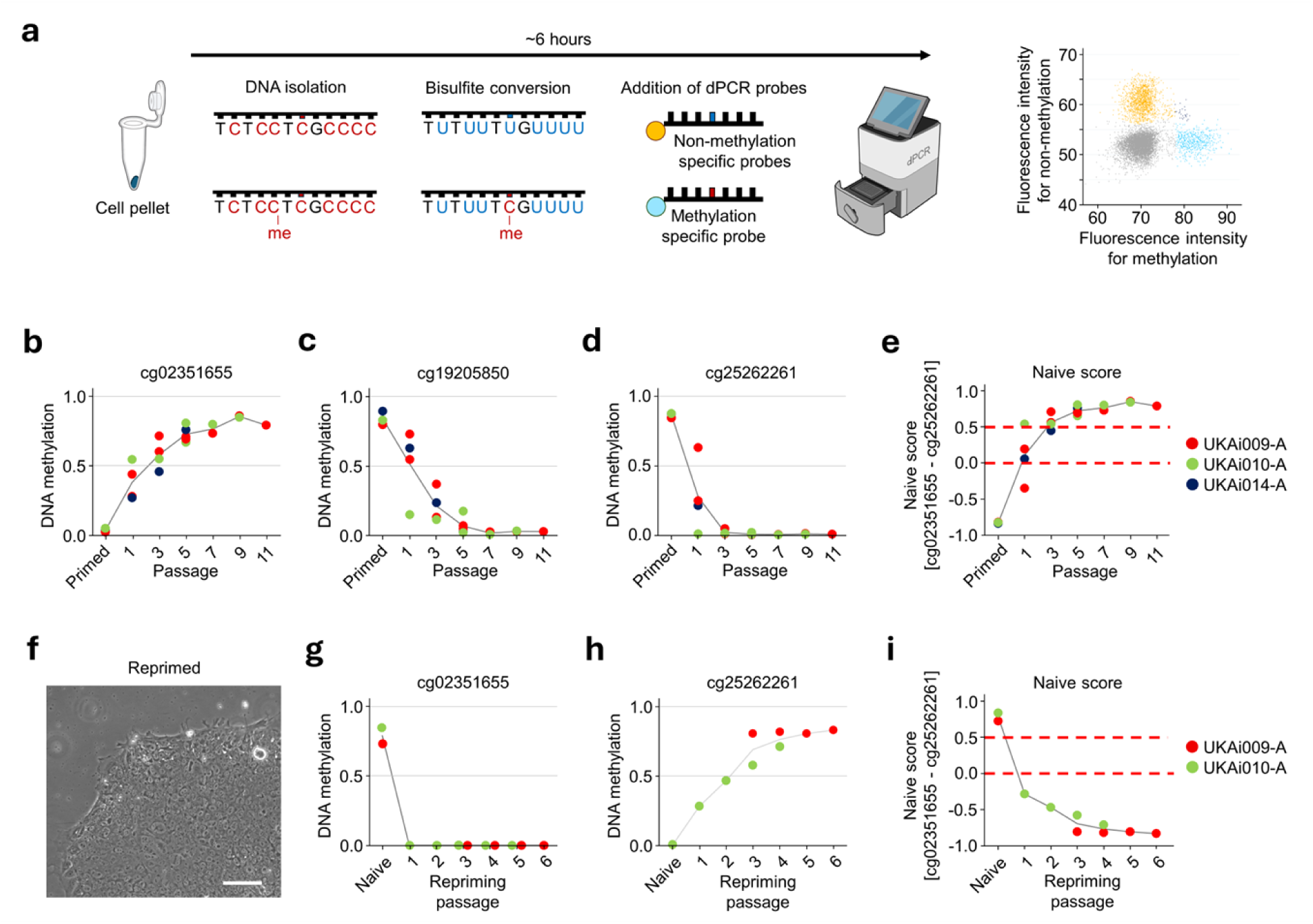
Analysis of naive-score with digital PCR. **(a)** Schematic presentation for targeted DNAm analysis of individual CpGs by dPCR. Non-methylated cytosines are bisulfite converted into uracil which will be PCR amplified as thymine while methylated cytosines are protected. Different fluorescently labeled probes for methylated and non-methylated strands can be detected and Poisson distribution is used to determine the DNAm levels. **(b-d)** DNAm analysis at the three candidate CpGs by dPCR revealed rapid DNAm changes during primed-to-naive conversion within two to five passages in FINO culture conditions. **(e)** The naive-score indicates that within three passages the FINO-treated cells reach the naive threshold. **(f)** Morphology of iPSCs upon repriming of FINO-treated cells for four passages in primed culture conditions. Scale bar = 100 µm. **(g-i)** DNAm levels at the two relevant CpGs and the naive score transition to primed state within few passages or repriming

Next, we analyzed if our FINO-treated naive cells could be reprimed upon transition back into primed culture medium. Within days, the cells showed typical phenotypic changes and after four passages they acquired homogeneous morphology of primed iPSC colonies (Figure 5f). Notably, digital PCR analysis of the naive-score also reflected the efficient epigenetic change to primed state within this time frame (Figure 5g-i). Thus, targeted analysis of our epigenetic signature by digital PCR provides a quantitative measure for the dynamic changes of primed-to-naive transition and during repriming.

## Discussion

Many culture protocols have been established for primed-to-naive transition of human PSCs, which may lead to a spectrum of naive states (39). In our study, we integrated components from various media to create a feeder-free and normoxic workflow. Most naive induction protocols require low oxygen of 1 to 5%, as hypoxia was shown to enhance reprogramming efficiency into iPSCs (43), and contributes to the regulation of differentiation in stem cells (44). Additionally, HIF-1α knockout inhibits the transition from naive to primed human embryonic stem cells (45). Recently, the addition of dibenzazepine (YO-01027) and a DOT1L inhibitor have been shown to facilitate maintenance of naive-state under non-hypoxic conditions (22), and this was also observed in our FINO medium. Furthermore, we demonstrated translocation of HIF-1α to the nucleus in FINO conditions, a phenomenon typically associated with hypoxic environments (44). FINO medium also eliminated the need for a supportive feeder layer. Thus, our FINO protocol can significantly simplify the application of the primed-to-naive transition.

While our FINO-treated cells exhibited classical hallmarks of naiveness, our analysis highlighted the necessity for developing robust biomarkers to directly compare with naive-like cell preparations generated across different laboratories. Global hypomethylation or even global demethylation is considered as a hallmark characteristic of primed-to-naive conversion (39, 46). However, our integrative analysis of ten DNAm datasets reveals that while numerous genomic regions exhibit hypomethylation, others remain stable or even gain methylation during the primed-to-naive conversion. The observation that certain genomic sites acquire DNA methylation is particularly intriguing and has so far received limited attention. This finding underscores that primed-to-naive conversion encompasses more than just global hypomethylation; it is associated with nuanced alterations within the methylation landscape.

Based on these findings, we developed an epigenetic signature to differentiate between primed and naive pluripotent cells. DNAm patterns are particularly well-suited for such classifications, as epigenetic modifications play a crucial role in cell fate decisions and can be analyzed quantitatively at single-base resolution (32). The use of DNA as input material provides practical advantages over cell-based assays (such as immunofluorescence or flow cytometry) or RNA-based methods (like gene expression analysis) since DNA is relatively stable, even at room temperature, which greatly simplifies bulk processing and transportation to central laboratories or service providers. Our naive-score is based solely on two CpG sites whose methylation states change reciprocally during the transition from primed to naive states. Most likely, these DNAm changes are not of functional relevance, *per se* – they rather serve as biomarkers indicative of broader genome-wide reorganization of the methylome, which appears to be orchestrated within complex epigenetic networks (47, 48).

A two-CpG-signature provides a trade-off between applicability and specificity. Targeted methylation read-outs based on individual CpGs have proven diagnostic power in tumor classification (49), blood-cell deconvolution (50), and authentication of pluripotent cultures via the Epi-Pluri-Score (51) or PluripotencyScreen (37). An important advantage of such signatures is their cost-effectiveness in measurement without reliance on specific profiling techniques or intricate bioinformatics tools (32). In this study, we describe a protocol for determining the naive-score via digital PCR, which can be pursued within six hours. This analysis clearly revealed the stepwise methylation changes in iPSCs during FINO treatment, highlighting its utility for routine monitoring, protocol optimization, and high-throughput screening applications. Alternatively, the naive-score might also be addressed by other methods for DNAm analysis, such as pyrosequencing or nanopore sequencing (52).

The multidimensional scaling analysis of global DNAm patterns, along with the naive-score, indicates that several published naive culture conditions do not fully replicate the naive DNAm state and tend to remain in an intermediate state (datasets 1-5). In fact, there appears to be a spectrum of naive states during early embryonic development, ranging from pre-implantation inner cell mass cells through a proposed formative state to the post-implantation primed epiblast (2). The naive-score may reflect this continuum of pluripotency in human stem cell culture, with 2iL/NHSM representing an intermediate naive state and FINO/FINE/t2iLGö achieving a more complete reset (39). It is also possible that these intermediate naive cell preparations consist of mixtures of both primed and naive subsets. Regardless, the naive-score offers a quantitative measure for benchmarking various naive culture protocols.

## Limitations

Our FINO protocol induced a naive phenotype in at least four out of six iPSC lines tested and it will be important to better understand whether some lines may be resistant to this conversion. While the FINO-treated cells exhibited many hallmarks of naive pluripotency, we could not directly compare them with all the other cell preparations or conduct assays such as chimera formation. It is also crucial to monitor extended culture under FINO conditions, as the multiple pathway inhibitors used may have unknown off-target effects. On the other hand, it would be intriguing to investigate whether the primed-to-naive conversion effectively eliminates all epigenetic traces from the tissue of origin and whether transient treatments could functionally correct iPSCs to restore unbiased multilineage differentiation potential (14). Our integrative DNAm analysis across various studies indicates that there exists a continuum of naive-associated DNAm changes. Single-cell transcriptomic analysis can reconstruct such trajectories (41), whereas single-cell DNAm profiling remains challenging and costly at present. Nevertheless, such single cell analysis profiling could provide deeper insights into the heterogeneity of naive and primed subsets, as well as potentially identify defined intermediate states. Lastly, due to ethical constraints, we are unable to benchmark our naive-score against human pre-implantation epiblasts.

## Conclusion

The FINO medium offers a straightforward approach to achieving naive human pluripotent stem cells by eliminating the need for xenogeneic feeders and hypoxia chambers. It addresses the significant technical and economic challenges associated with routine naive stem cell research. Nonetheless, there is still a pressing need for systematic optimization of these culture conditions and to validate successful conversion with a standardized assay (39). The naive-score provides a rapid and cost-effective tool for quantitative assessment of the pluripotent state, facilitating routine in-process monitoring and high-throughput screening.

## Methods

### Human pluripotent stem-cell lines and maintenance

For the data presented in this manuscript we used three human iPSC lines, registered at human pluripotency stem cell registry (hPSCreg)(53): UKAi009-A, UKAi010-A, and UKAi014-A, which were derived from mesenchymal stromal cells (54). All samples were obtained after informed and written consent in accordance with the Declaration of Helsinki and the research was approved by the local ethics committee of RWTH Aachen University (EK206/09 and EK128/9). The iPSC lines were cultured and expanded under primed conditions on growth factor–reduced Matrigel (Corning) in iPS-Brew (Miltenyi Biotec) with daily medium changes and incubated at 37°C, 5% CO₂, and atmospheric O₂. Routine passaging (1:4-1:10 ratio) was performed when cells reached 70–80% confluency with 0.5 µM EDTA.

### Primed-to-naive conversion

For primed-to-naive conversion, we established FINO medium (Feeder-Independent, Normal Oxygen), which is based on components of the feeder-independent FINE formulation (Szczerbinska et al., 2019) and the non-hypoxia, LIF-dependent NHLD medium (Isono et al., 2020). FINO medium consists of N2B27 base (1:1 DMEM/F-12 and Neurobasal, 1% N2, 2% B27, 2 mM L-glutamine, 1% non-essential amino acids, 100 µM 2-mercaptoethanol; all ThermoFisher Scientific), 62.5 ng/mL BSA (Sigma) supplemented with: 1 µM PD0325901, 0.1 µM Dasatinib, 0.1 µM AZD5438, 0.1 µM SB590885, 10 µM Y-27632, 3 µM Pinometostat, 2 µM YO-01027 (all MedChemExpress), 20 ng/mL hLIF (ThermoFisher Scientific), 20 ng/mL Activin A (StemCell Technologies), 8 ng/mL FGF2 (Peprotech), 100 µM L-ascorbic acid 2-phosphate sesquimagnesium salt hydrate (Merck). For primed-to-naive conversion, primed cells were dissociated into single cells with Accutase (PanBiotech), resuspended in iPS-Brew + 10 µM Rho-associated protein kinase (ROCK) inhibitor Y-27632 (MedChemExpress) and plated on Matrigel-coated plates (10 000 cells/cm²). After 24 h Y-27632 was removed from the medium and after another 24 h medium was switched to FINO medium, and continuously changed every 24 h. Cells were passaged (1:2-1:6 ratio) when they reached 70-80% using Accutase.

### Immunofluorescence

Cells were fixed in 4% paraformaldehyde (15 min), permeabilized and blocked with 0.1% Triton X-100 and 1% BSA (60 min). Primary antibodies: KLF17 (Merck, 1:300), HIF-1α (Clone: MOP1, BD Biosciences, 1:200), SUSD2::PE (Clone: W5C5, Biolegend, 1:80), were incubated for 1h at room temperature (RT). Secondary antibodies anti-mouse::Alexa Fluor 488 (ThermoFisher Scientific, 1:125) or Anti-rabbit::Alexa Fluor 488 (ThermoFisher Scientific, 1:125) were applied for 30 min at RT. Nuclei were stained with DAPI (10 ng/mL) for 15 min at RT. Images were captured on a Zeiss Axio Observer 5 microscope and processed with Zen v3.10.

### Flow cytometry

Single-cell suspensions were fixed in 4% PFA (15 min) and incubated for 30 min at 4 °C with SUSD2::APC (Clone: W5C5, BioLegend, 1:80), CD75::PE (Clone: LN1, BD Bioscience, 555654, 1:80), or CD90::APC (Clone: 5E10, Biolegend, 1:80). Data were acquired on a BD Canto II and analyzed with FlowJo v10.9.

### DNA methylation profiling

The DNAm profiles were analyzed in primed iPSCs and after seven or ten passages of FINO-treatment in three independent replicas of UKAi-009-A. Genomic DNA was isolated with the AllPrep Kit (Qiagen). DNAm profiles were analyzed with Infinium Methylation EPIC v2.0 BeadChip technology at LIFE & BRAIN GmbH (Bonn, Germany). In addition, we used Illumina BeadChip datasets (450k and EPIC v1.0) from 10 published datasets (Supplementary Table S1), which were downloaded from NCBÍs Gene Expression Omnibus (GEO): For the datasets GSE102031, GSE95531, GSE208299, GSE179473, GSE128125, GSE128126, and GSE224697 the IDAT files were pre-processed with minfi (55) (v1.54.1) in R (v4.4.2). Low-quality samples were removed (threshold: sum of the medians of the methylated and unmethylated channels <20), and the remaining samples were normalized with ssNoob (56) (Sesame v1.26.0). For samples where no IDAT files were available (GSE57318, GSE65214, and GSE288982), we used already existing beta values or generated the beta values from the signal intensities. CpG sites on XY chromosomes were not considered, unless stated otherwise. Furthermore, we did not consider non-CG probes and SNP-associated CpGs for further analysis. For integrative analysis across the different studies, we only focused on CpGs that were represented on every dataset. The association of CpGs to promotor regions was defined by TSS1500 and TSS200 from the Illumina annotation file. The limma R package was used for calculation of Benjamini-Hochberg adjusted p values and the MDS plots. Relevant DNAm changes were defined as showing at least 20% difference in mean DNAm values and an adjusted p value < 0.05. The R packages ggplot2 (v3.5.2) and ggrepel (v0.9.6) were used for graphical presentation.

### Selection of epigenetic biomarkers

The selection of candidate marker CpGs was conducted using the R package CimpleG (v0.0.5.9028) (36) to identify CpG sites that exhibit substantial differences in mean DNAm values while demonstrating low variance within the groups. In addition to the above mentioned datasets for naive and primed pluripotent stem cells, we also considered DNAm profiles of early differentiation towards endo-, meso-, and ectoderm (GSE207119) (37), to exclude that our signature might be confounded by early differentiation events. Based on this we identified three CpGs that can even individually reveal good distinction between naive and pluripotent cells: cg02351655, cg19205850, and cg25262261. To further ease applicability, we subsequently utilized two of these CpGs for the naive-score (naive-score = DNAm [cg02351655] – DNAm [cg25262261]) and the threshold was arbitrarily set at 0. To further benchmark this score, we used 275 samples of primed stem cells from the HipSci iPSC bank database (https://www.hipsci.org/#/; accessed on May 2, 2025) (38), as well as DNAm profiles generated in the course of reprogramming fibroblasts into iPSCs (GSE54848) (42).

### Gene expression analysis

RNA was isolated with the AllPrep Kit (Qiagen). For semiquantitative RT-PCR the RNA (500 ng) was converted to cDNA with the high-capacity cDNA RT kit (ThermoFisher Scientific) according to the manufacturers protocol. 10 ng of cDNA was mixed with iTaq SYBR master mix (Bio-Rad) and 0.5 µM of forward and reverse primer (Supplemental Table S2) and analyzed on a CFX Duet (Bio-Rad). Cycling: 95°C 30 s, 40 × (95°C 3 s, 50°C 20 s), melt curve from 65°C to 95°C in 0.5°C increments. Data were analyzed with Bio-Rad CFX Maestro 5.3.022.1030 software.

Bulk RNA sequencing was performed on primed and FINO-treated cells for each of UKAi009-A and UKAi010-A, each in two independent replicas by the West German Genome Center (Cologne, Germany) utilizing a NovaSeq 6000 sequencer configured for a 1 x 100 single-end format, resulting in an average of 10-20 million reads per sample. Following quality control with FastQC (v0.12.1) (57), reads were trimmed using Trim Galore! (v0.6.10) (58)and aligned to the human reference genome (hg38) employing STAR alignment software (v2.7.11b) (59). A count table of the mapped reads was generated using Salmon software (v1.10.3) (60). Transcripts with fewer than 10 reads were excluded from further analysis.

Differential expression analysis was conducted using the DESeq2 package (v1.32.0) (61), which facilitated the logarithmic transformation of the data and subsequent exploratory analysis. Gene counts were normalized using a variance-stabilized transform (VST), and Principal Component Analysis (PCA) of the VST data was performed using the plotPCA function from the DESeq2 package. Visualizations were generated using ggplot2 (v3.3.6) (62). The DESeq2 package was used to perform differential expression analysis, where adjusted p-values were computed using the Benjamini–Hochberg method. Genes were considered differentially expressed if they had an adjusted p-value of less than 0.05 and a log2 fold change greater than or equal to 0.5 or less than or equal to-0.5. Gene annotations were incorporated into the resulting files using the biomaRT package (v2.48.3) (63). To explore the correlation between DNAm and gene expression data we utilized the Illumina BeadChip annotation by matching to Ensembl IDs, focusing on CpGs located within promoter regions, which were defined by TSS1500 and TSS200 from the Illumina annotation file. Gene set enrichment analysis (GSEA) utilized pre-ranked genes (based on the fold change in DESeq2) and performed 100,000 permutations. Functional enrichment analysis was performed using fgsea (v1.20.0). Adjusted p-values were calculated using the Benjamini–Hochberg method.

### DNA methylation analysis with digital PCR

Genomic DNA (500 ng) was bisulfite-converted using either the EZ DNA Methylation Kit (Zymo Research) or the EpiTect Fast Bisulfite Conversion kit (Qiagen). 25 ng of DNA was mixed with QIAcuity Probe PCR master mix (Qiagen), 0.8 µM forward and reverse primers and 0.4 µM probe primers (Supplemental table S3) and partitioned into 8.5k chambers on a QIAcuity, 5 plex machine (Qiagen). Cycling: 95°C 2 min, 40 × (95°C 15 s, 56°C 30 s); imaging was set at green: 300 ms, gain 6, yellow: 250 ms, gain 4. Data were analyzed with the QIAacuity Software Suite 3.1.0.0.

## Declarations

## Acknowledgements

This work was supported by the Flow Cytometry Facility, a core facility of the Interdisciplinary Center for Clinical Research (IZKF) Aachen within the Faculty of Medicine at RWTH Aachen University and by the German Research Foundation (DFG), project ID 439895892. Funding of this work was supported by the Deutsche Forschungsgemeinschaft (DFG: 363055819/GRK2415; WA1706-12-2/CRU344/417911533; WA1706-14-1/458369830; WA1706-17-1/561150360; and SFB 1506-1/450627322), by the José Carreras Foundation (DJCLS 03 R/2024), and particularly by the Federal Ministry of Education and Research (VIP+: PluripotencyScreen; 03VP11580).

## Author Contributions

I.O.S.: Conceptualization, Methodology, Formal analysis, Investigation, Data Curation, Writing of original draft, Visualization, Supervision. T.A.: Investigation, Validation. R.K.: Formal analysis, Data Curation, Visualization. K.Z.: Contribution to writing. W.W.: Conceptualization, Formal analysis, Resources, Writing of original draft, Supervision, Funding acquisition.

