## Supplemental Figures and Tables for "Primed-to-naive conversion of pluripotent stem cells can be tracked by specific DNA methylation changes for optimized culture conditions"

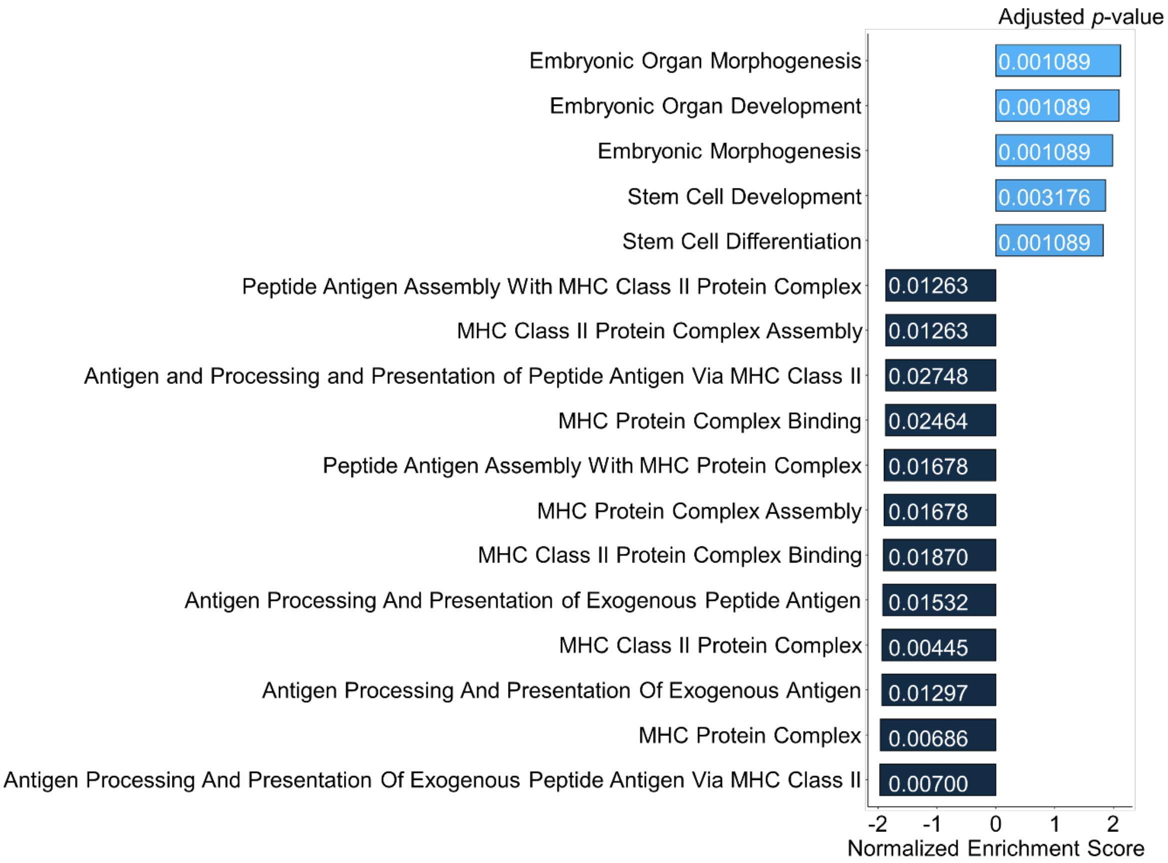


**Figure S1. Gene Ontology analysis of differential gene expression upon FINO-treatment.** The differentially expressed genes in iPSCs upon treatment with the feeder-independent, normoxic protocol (FINO) was further analyzed by Gene Ontology classification. The 1024 up-regulated and 624 down-regulated genes (adjusted *p*-value < 0.05, Log2 fold change > 1) were clearly enriched in embryonic and stem cell related categories, and major histocompatibility complex (MHC) related genes, respectively.

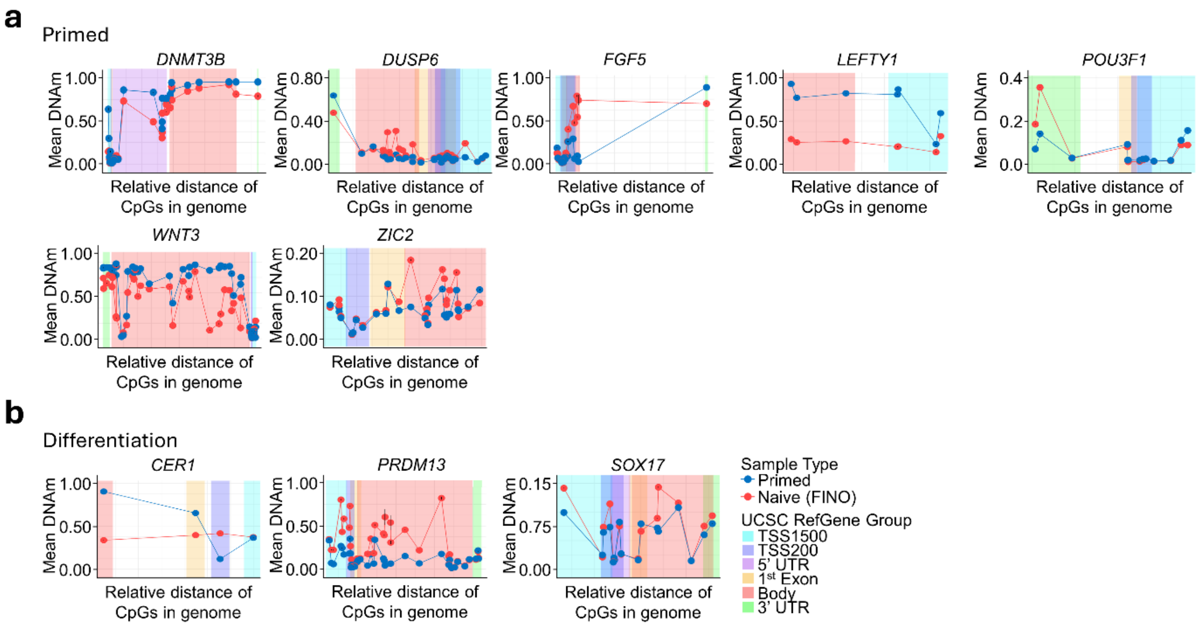


**Figure S2. DNA methylation changes in genes related to primed state or differentiation.**
This figure depicts DNA methylation patterns in representative genes that are up-regulated **a)** in primed state (*DNMT3B, DUSP6, FGF5, LEFTY1, POU3F1, WNT3,* and *ZIC2*), or **b)** during differentiation (*CER1, PRDM13*, and *SOX17*). While many of the representative primed-associated genes also reveal hypomethylation in the promotor regions, in analogy to the naive-associated genes (Figure 1g), there were also some genes with opposite effects, such as *DUSP6*, *POU3F1*, *PRDM13,* and *SOX17*).

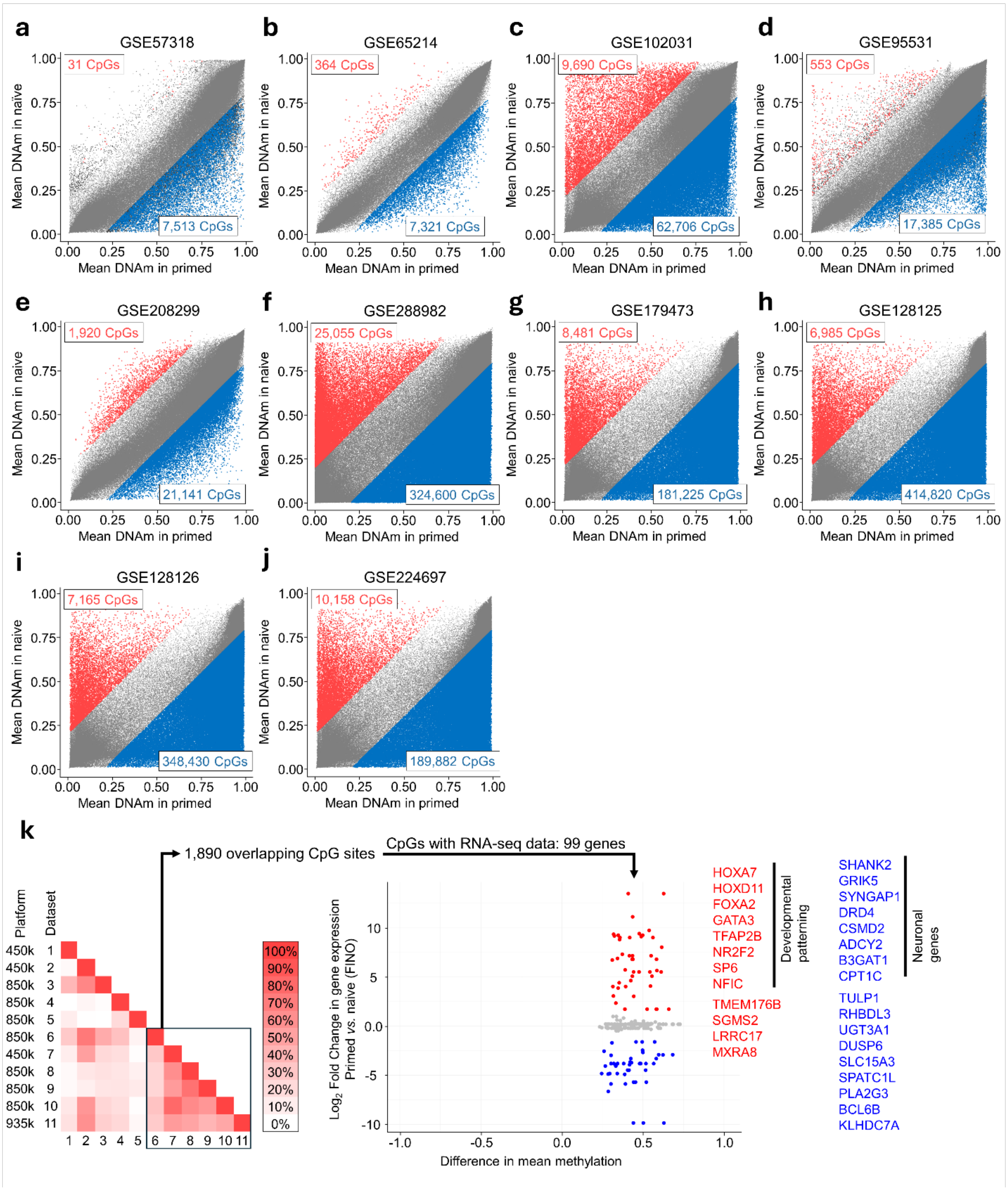


**Figure S3. DNA methylation changes in primed-to-naive conversion in individual datasets.** **a-j)** Scatter plots of differential methylated CpGs in datasets 1-10 (adjusted *p*-value: < 0.05, diff. mean DNAm > 0.2). The GEO IDs are indicated for each dataset as well as the number of significantly hyper- (red) and hypomethylated CpGs (blue) in naive *versus* primed state. **k)** To further elucidate the gene-association of CpGs that become hypermethylated in primed-to-naive conversion, we focused on the datasets 6-11, which have a higher fraction of overlapping CpGs (left pane). 1,890 CpGs were overlapping hypermethylated in datasets 6-11 (adjusted *p*-value < 0.05, mean diff. DNAm > 0.2), which were associated with 99 genes identified in our RNA-seq analysis. When we focused on differentially expressed genes within these 99, 8 out of 12 up-regulated genes in naive state were associated with developmental patterning while 8 out of 17 down-regulated genes were associated with neuronal genes (adjusted p-value < 0.05, fold change > 2).

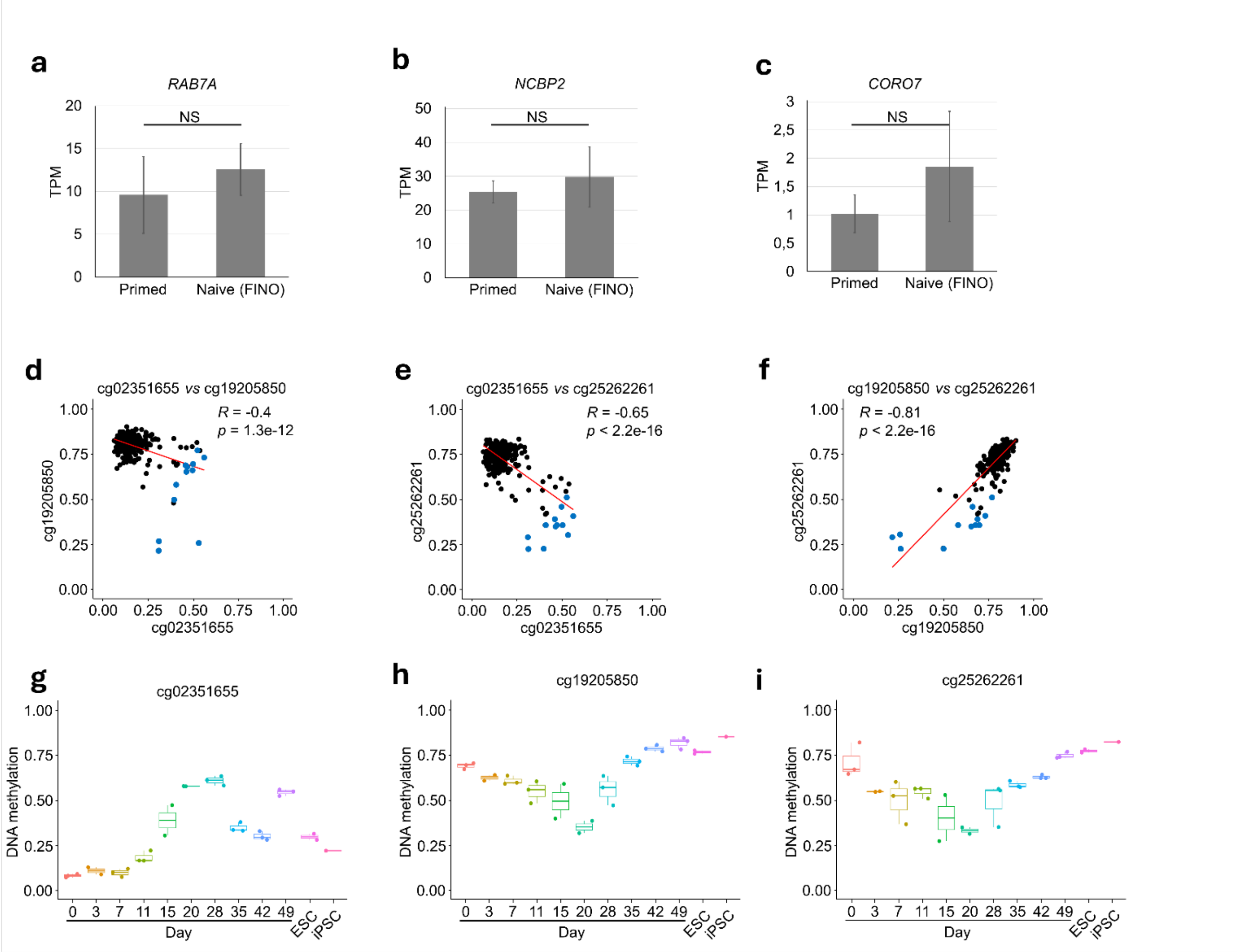


**Figure S4. Further analysis of CpGs of the naive-score.**
**a-c)** Gene expression changes were analyzed in the three genes corresponding to the three candidate CpGs for naive *versus* primed comparison (cg02351655: *CORO7*; cg19205850: *RAB7A*; and cg25262261: *NCBP2*). Bar plots depict transcripts per million (TPM) in RNA-seq analysis (NS = $p$-value > 0.05 via Student's t-test; n = 4). **d-f)** To further explore why few samples of the HipSci stem cell bank were above thresholds of naive state for individual CpGs, we performed scatterplot analysis to determine if these aberrations were consistent between our CpGs. In fact, all CpGs showed a clear correlation, indicating that the deviations are not due to technical noise, but rather a more naive-like DNAm landscape in these samples (highlighted in blue; Pearson correlation coefficient and $p$-value are indicated). **g-i)** We further analyzed the dynamics of the three candidate CpGs during reprogramming of somatic cells into iPSCs using the dataset GSE54848. Notably, all three CpGs indicate the same transient change toward naive-like DNAm patterns around day 20.

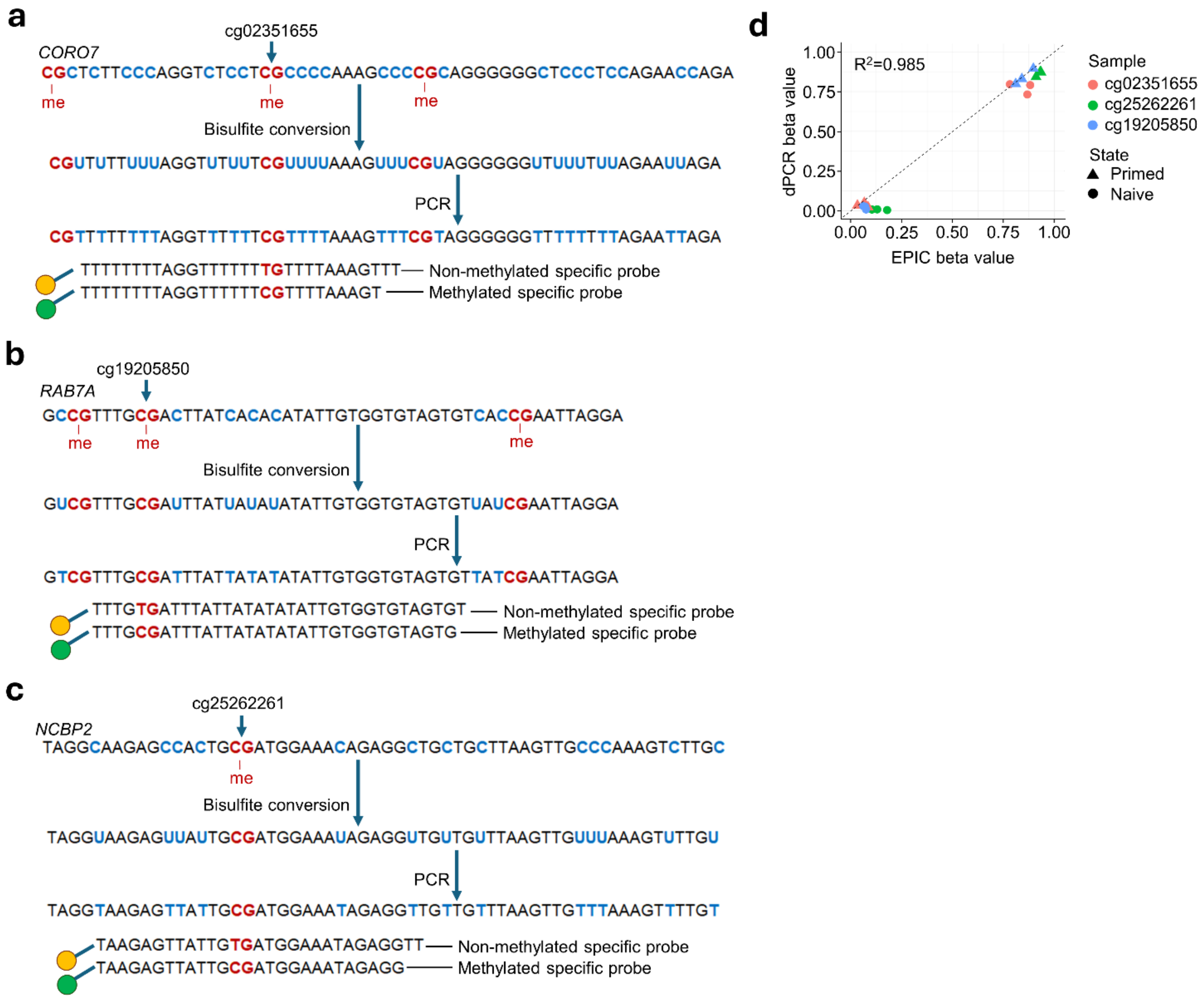


**Figure S5. Digital PCR assays to investigate primed to naive transition.**
**a-c)** Schematic presentation of the selected CpGs and their probe designs to investigate DNA methylation by dPCR. The probes are labelled either with HEX (yellow) and FAM (green) fluorophores. **d)** Scatter plots to compare the measured DNAm levels in the Illumina BeadChip data and dPCR. The different colors depict the three candidate CpGs and the symbols represent primed and naive state. The coefficient of determination ($R^2$) demonstrates high correlation between DNAm levels determined with the two different methods.

### Supplemental Tables

**Supplemental Table S1: Datasets used in this study.**

| Dataset | Acc. Nr. | Platform | Species | Cell types | Cell lines | Datatype |
|---|---|---|---|---|---|---|
| 1 | GSE57318 | 450k, GPL13534 | Human | Primed and naive hiPSC | 2 | Signals |
| 2 | GSE65214 | 450k, GPL13534 | Human | Primed and naive hiPSC | 5 | Signal Intensities |
| 3 | GSE102031 | EPIC, GPL21145 | Human | Primed and naive hiPSC | 2 | IDAT |
| 4 | GSE95531 | EPIC, GPL21145 | Human | Primed and naive hiPSC | 1 | IDAT |
| 5 | GSE208299 | EPIC, GPL23976 | Human | Primed and naive hiPSC | 4 | IDAT |
| 6 | GSE288982 | EPIC, GPL21145 | Human | Primed and naive hiPSC | 1 | Normalised Beta |
| 7 | GSE179473 | 450k, GPL13534, EPIC, GPL21145 | Human | Primed and naive hiPSC | 3 | IDAT |
| 8 | GSE128125 | EPIC, GPL21145 | Human | Primed and naive hiPSC | 1 | IDAT |
| 9 | GSE128126 | EPIC, GPL21145 | Human | Primed and naive hiPSC | 1 | IDAT |
| 10 | GSE224697 | 450k, GPL13534, EPIC, GPL21145 | Human | Primed and naive hiPSC | | IDAT |
| 11 | Own | EPICv2 | Human | Primed and naive hiPSC | 1 | IDAT |

**Supplemental Table S2: Primer sequences for RT-PCR.**

| Gene target | Forward/Reverse | 5' → 3' |
|---|---|---|
| *DPPA3* | Forward | GGACGATTCAGGGGCCTCTCAA |
| *DPPA3* | Reverse | CCCTGAGGACTGCTGCTCCTAC |
| *DPPA5* | Forward | ACAAGCTCCGGACCAAGTGGAT |
| *DPPA5* | Reverse | GCAAGTTTGAGCATCCCTCGCT |
| *KLF5* | Forward | ATTTACCCACCACCCTGCCAGT |
| *KLF5* | Reverse | TGTGCAACCAGGGTAATCGCAG |
| *TFCP2L1* | Forward | ACCTGTCTGTGTACCACGCCAT |
| *TFCP2L1* | Reverse | CCATCTCGTTGCTCACCACCAC |
| *DNMT3L* | Forward | CCCGGGACAGGCAGAGATCTTA |
| *DNMT3L* | Reverse | TGTCCTTACATGGGGCGCAGAT |
| *ZIC2* | Forward | AAACTGGTCAACCACATCCGCG |
| *ZIC2* | Reverse | CTGGAACGGCTTCTCCCCTGTG |
| *B3GAT1* | Forward | GCCAACTGCACCAAGATCCTGG |
| *B3GAT1* | Reverse | GAGGCTCAGATCTCCACCGAGG |
| *TBP* | Forward | TGCACAGGAGCCAAGAGTGAAGA |
| *TBP* | Reverse | TTCACATCACAGCTCCCCACCA |

**Supplemental Table S3: Primer and probe sequences for digital PCR.**

| CpG ID | Forward/Reverse/Probe | 5' → 3' |
|---|---|---|
| cg02351655 | Forward | TTAGTTAGTAGTTGGAAGGG |
| cg02351655 | Reverse | TACTATCCAAAAAACTCCCA |
| cg02351655 | Unmethylated Probe | HEX-TTTTTTTTAGGTTTTTTTGTTTTAAAGTTT-MBQ-1 |
| cg02351655 | Methylation Probe | FAM-TTTTTTTTAGGTTTTTTCGTTTTAAAGT-MBQ-1 |
| cg19205850 | Forward | TTGATGTTATTTGAAAGTTTTTT |
| cg19205850 | Reverse | TCAAACCTCAATTAAATACCCT |
| cg19205850 | Unmethylated Probe | HEX-TTTGTGATTTATTATATATATTGTGGTGTAGTGT-MBQ-1 |
| cg19205850 | Methylation Probe | FAM-TTTGCGATTTATTATATATATTGTGGTGTAGTG-MBQ-1 |
| cg25262261 | Forward | AGTTTTTTAAAGTGTTGGGATT |
| cg25262261 | Reverse | CACATCCCACTCTACAAAACT |
| cg25262261 | Unmethylated Probe | HEX-TAAGAGTTATTGTGATGGAAATAGAGGTT-MBQ-1 |
| cg25262261 | Methylation Probe | FAM-TAAGAGTTATTGCGATGGAAATAGAGG-MBQ-1 |
